# SEX-SPECIFIC IMPACTS OF EARLY LIFE SLEEP DISRUPTION: ETHANOL SEEKING, SOCIAL INTERACTION, AND ANXIETY ARE DIFFERENTIALLY ALTERED IN ADOLESCENT PRAIRIE VOLES

**DOI:** 10.1101/2024.01.03.574112

**Authors:** Darren E. Ginder, Carolyn E. Tinsley, Mara E. Kaiser, Miranda M. Lim

## Abstract

Early life sleep is important for neuronal development and maturation. Using the highly social prairie vole rodent model, we have previously reported that early-life sleep disruption (ELSD) during the pre-weaning period postnatal day (P)14 to 21 results in adult interference with social bonding and increases ethanol consumption following a stressor. Furthermore, we have reported increased parvalbumin expression and reduced glutamatergic neurotransmission in cortical regions in adult prairie voles that experienced this paradigm. To understand the impact of ELSD on the lifespan, examination of an earlier time in life is necessary. Thus, the aim of the present study was to examine the behavioral outcomes of ELSD on adolescent prairie voles. Here we hypothesized that anxiety and reward related behaviors, as measured by light/dark box, 2-bottle choice and social interactions, would be negatively impacted by ELSD in adolescent male and female prairie voles. Male ELSD voles were no different from control voles in measures of anxiety and ethanol preference or consumption, but affiliative social interactions were significantly reduced. ELSD differentially impacted female prairie voles, with increased anxiety-like behavior and reductions in ethanol consumption compared to Controls, but no impact on ethanol preference or social interactions. Together, these results suggest both male and female prairie voles experience differential changes to reward seeking behaviors, but only female prairie voles showed increases in anxiety-like behavior. These results further suggest that early-life sleep is critically important for neurotypical behaviors in adolescence, a time where reward-seeking and risky behaviors are adaptive for learning and promoting survival.

## INTRODUCTION

Early life sleep is important neuronal development (see Alrousan et al., 2022 for a review). Importantly, early life sleep is skewed heavily towards rapid eye movement (REM) sleep, with large portions of total sleep time spent in REM in newborn humans (Roffwarg et al., 1966; Louis et al., 1997). Non-REM sleep has been highly correlated with neuronal development. Using electroencephalogram (EEG), slow-wave activity during sleep in early life has been shown to modulate brain plasticity (Wilhelm et al., 2014) and cortical maturation has been shown to be strongly correlated with slow-wave sleep activity (Buchmann et al., 2010; Kurth et al., 2010). Furthermore, the changes seen in slow-wave activity in adolescence have been thought to be related to adolescent synaptic pruning, as cortical density has been shown to reduce during this time (Shaw et al., 2008). This change in synaptic density does seem to be regulated by sleep, as sleep deprivation results in alterations to the synaptic proteome. Juvenile mice that experienced sleep deprivation show differences in their synapse proteome expression in adolescence and adulthood (Gay et al., 2023) compared to non-sleep deprived controls. REM sleep is also critically important to the developing brain. REM sleep deprivation of rat pups resulted in significant reductions to the excitability of the prefrontal cortex (PFC), with reduced expression of subunits for AMPA and NMDA receptors (Atrooz et al., 2019), suggesting REM sleep is important for neuronal plasticity. This is the case, as REM sleep has been shown to promote cortical plasticity during neonatal neurodevelopment in cats (Dumoulin Bridi et al., 2015). The need for REM sleep in early life in humans is theorized to be the same, with REM sleep being an important mechanism to foster brain development (Roffwarg et al., 1966).

Autism spectrum disorder (ASD) may be influenced by disruptions in early life sleep. Children diagnosed with ASD show increased sleep disturbances (Krakowiak et al., 2008) and shorter sleep duration (Veatch et al., 2017). These differences in sleep likely result in skewed developmental trajectories, resulting in the life-long cognitive and social deficits (Hajri et al., 2022). The disruption of sleep in early life may increase vulnerability of individuals towards the development of ASD. Genetically susceptible mice (*Shank3*) have shown increased vulnerability towards ASD-related pheno-conversion following early life sleep disruption (ELSD; Lord et al., 2022), providing further support for the importance of early life sleep in neurotypical development. As ASD diagnoses rely on deficits in social interactions (American Psychiatric Association, 2013), examination of social behaviors is important for determining the ASD-relevent impacts of ELSD. To that end, the use of a highly social animal model is necessary. Prairie voles (*microtus ochrogaster)* are a wild caught rodent that serve as an excellent model of human social behavior, due to similarities in pair-bonding and social monogamy that are displayed in both humans and prairie voles (Carter et al., 1995). Both humans and prairie voles have shown that pair-bonded individuals live longer, compared to their single counterparts (House et al., 1988) and that both species exhibit similar sociosexual attachments to their pair-bonded partner (Young et al., 2011). Previously our laboratory has reported that ELSD heavily influences pair bonding behavior in adult prairie voles. Adult male prairie voles that experienced ELSD showed reduced preference for their female partner during a partner preference test as evidenced by reduced side by side social huddling (Jones et al., 2019). Furthermore, adult voles that underwent ELSD showed increased ethanol consumption following acute, inescapable foot shock (Jones et al., 2020) despite no differences in baseline ethanol consumption compared to non-sleep disrupted Controls. Finally, ELSD significantly reduced the amount of time adult male prairie voles spent with a novel object (Jones et al., 2019). As activity in the nucleus accumbens is involved in novel object interactions (Bevins et al., 2002; Bevins & Besheer, 2005), the changes to novel object interaction reported by Jones et al. (2019) suggest alterations to reward circuitry. Together, these data imply that ELSD results in longitudinal impacts in brain and behavior. With this long-term impact of ELSD in mind, there may be some change in adolescent behavior as well.

Adolescence is an important timepoint for learning and neurodevelopment (Spear, 2013). Importantly, adolescence is a timepoint where reward seeking and risky behaviors are seen to increase (Galvan, 2010). This may be related to changes in the nucleus accumbens (NAc), both volume (Hammerslag & Gulley, 2016) and dopamine receptors D1 and D2 (Andersen et al., 1997) show increases and decreases during this time. This adolescent increase in reward sensitivity and risky behavior has been often viewed as a negative trait with the perspective of reducing potential risk to adolescent children (Telzer, 2016). More recent theories suggest the opposite, that this time of increased reward sensitivity and risky behavior is adaptive for learning and promoting survival when caregiver safety is no longer available (Spear, 2000; Wahlstrom et al., 2010; Telzer, 2016).

Individuals diagnosed with ASD have shown to experience reduced reward sensitivity (Kohls et al., 2013). This may be related to dopaminergic dysfunction in the NAc (Paval, 2017). In 18 males diagnosed with ASD, fMRI does reveal reduced activation of the NAc during a go/no-go task (Kohls et al., 2013), highlighting dopaminergic dysfunction is the case in these individuals with ASD. Further evidence for dopaminergic dysfunction is provided by Baumester et al. (2023), where fMRI was performed on 212 individuals diagnosed with ASD during a social and monetary reward task. Imaging revealed that ASD individuals experienced hypoactivation of the ventral striatum (*i.e.* NAc) during both social and monetary reward tasks (Baumester et al., 2023). Hypoactivation of the NAc highlights that ASD individuals do not experience reward sensitivity in the same manner as their neurotypical counterparts. As previously shown, ELSD enhances ethanol consumption following a stressor (Jones et al., 2020) as well as impairing social bonding (Jones et al., 2019) in prairie voles. Expanding the age of observation is essential towards understanding the impact of ELSD on neurodevelopment.

With these considerations, the present study aimed to examine reward seeking and risky behavior in adolescent prairie voles through two-bottle choice (2BC), with voles choosing between ethanol, at increasing concentrations, and plain tap water. We propose that adolescent prairie voles that experienced ELSD exhibit reductions in ethanol preference and consumption. Additionally, due to alterations in social behavior in adult prairie voles that experienced ELSD (Jones et al., 2019), we expect to see deficits in social interactions between siblings in adolescent ELSD voles. Furthermore, due to adult voles that experienced ELSD showing an inability to cope with stressful events (Jones et al., 2020) we predict adolescent prairie voles show heightened anxiety.

## METHODS

### Animals

Prairie voles were housed in rooms that were controlled for temperature and humidity and had a 14:10 light/dark cycle (lights on at 0700h). Food and water access was *ad libitum* and cotton nestlets were provided for nesting material with weekly cage changes. Weaning occurred on P21, when animals were removed from parents and housed in same sex sibling pairs (2 per cage). Following weaning, females were housed in a separate colony room from males, due to female prairie voles being induced ovulators, and males and females were tested separately. As such, we do not directly compare males and females or use sex as a grouping variable. It was presumed that all females were in anestrus throughout the experiment due to the lack of male presence in the female only colony room. Subjects in these experiments were sourced from 12 breeder pairs and 25 unique litters. Male and female prairie voles underwent either ELSD or control (CON) sleep conditions from P14 – P21 and were left undisturbed until adolescence (see Figure 1 for experimental timeline; ELSD/male = 20, ELSD/female = 18, CON/male = 18, CON/female = 16).

**Figure 1.**
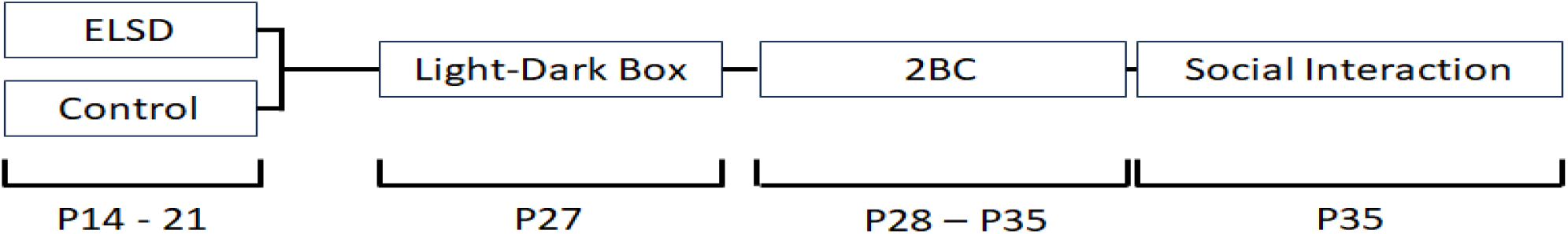
Experimental timeline

### Early Life Sleep Disruption (ELSD)

From P14-P21, voles underwent ELSD by placing home cages containing prairie vole litters on a laboratory orbital shaker (Jones et al., 2019, Jones et al., 2020, Jones et al., 2021, Jones et al 2022). Litters were maintained on shakers continuously from P14 to P21, when litters were weaned. Auditory disruption was minimized by locking cage cards in place and cage flooding was eliminated by replacing water bottles with hydrogel. Orbital shakers were connected to an automatic timer which gently agitated (110RPM) home cages for 10 s every 110 s (see Jones et al., 2019 and Jones et al., 2020 for additional details). Control (CON) voles were placed in the same room as ELSD voles from P14 to 21, but sleep was not disrupted. While we have previously shown that this method of sleep disruption significantly reduces REM sleep and fragments non-REM (NREM) sleep compared to controls, this paradigm does not impact parental care, pup weight, or serum corticosterone levels in pups, suggesting that this method results in no developmental impact of stress (Jones et al., 2019).

### Light-Dark Box

On P27, prior to exposure to ethanol, prairie voles underwent a 10 minute light/dark box test to quantify anxiety-like behaviors. The light/dark box was a modified open field (60 cm x 60 cm x 30 cm; Omnitech Electronics, Columbus, OH) with a dark insert (30 cm x 30 cm x 20 cm) and a door opening (10 cm x 5cm) allowing the animals to freely cross from one side to the other. A lux level of 3000 was achieved in the light portion by supplementing room light with additional lamps. Voles were placed in the lit compartment facing the outer wall at the beginning of the test and allowed to ambulate freely. Automatic beam break data was recorded in both the light and dark zones.

### Two-Bottle Choice (2BC)

At P28, home cages had water bottles removed and replaced with two 25mL glass cylinder tubes affixed with sipper tubes – one filled with standard drinking water from an autoclaved water bottle and the other filled with increasing concentrations of ethanol (EtOH) in drinking water (v/v). EtOH concentration began at 3% EtOH and increased to 6% EtOH after two days. After two days of 6% EtOH, the concentration increased again to 10% EtOH for another four days (eight days of 2BC in total). This volume was chosen in order to provide *ad libitum* access to water or EtOH for the duration of the 2-bottle choice (2BC) test. In order to allow semisocial housing (*e.g.* allowing exposure to bedding of cage mate and limited physical contact) but restrict access to only one pair of drinking tubes, home cages were divided in half via chicken wire mesh (7 mm openings) (Anacker et al., 2014; Walcott & Ryabinin, 2017; and Jones et al., 2020). Bottles were read daily at 0800h, filled with fresh solution, and sides were alternated in order to avoid bias. To account for spillage, an empty cage with sipper tubes was placed on each rack: spillage was never more than 0.05mL. Total volume consumed from both bottles, amount of ethanol consumed (g/kg), as well as EtOH preference were all recorded daily. EtOH preference was calculated as a ratio of fluid consumed from the ethanol bottle divided by total fluid consumed from both bottles (EtOH and water).

### Social Interaction Test

P35 bottles were measured and removed. Sibling pairs were placed together in a clean home cage and observed for social behaviors for 10 minutes. The following behaviors were quantified for each pair of animals: autogrooming time (sec); huddling and allogrooming time (sec). Huddling and allogrooming time were grouped together into a single variable: affiliative behavior duration (sec). Affiliative behavior duration was used for analyses.

### Statistics

A repeated measures Analysis of Variance (ANOVA) was used to analyze 2BC and light/dark box data with condition (CON/ELSD) as between group factors and day (2BC) or lighting condition (light/dark box) as the within group factors. Social interaction data was analyzed via between-subject measures ANOVA and Pearson’s *r* analyses were conducted to determine if a relationship was present between ethanol consumption and social interactions. To further examine the differences based on sex, analyses were conducted on males and females separately. Statistical analyses were conducted with IBM SPSS Statistics (Version 27). Statistical significance was considered for *p* values ≤ .05 and effect size estimates are provided for all effects with *p* values ≤ .08.

## RESULTS

### Early-life sleep disruption increases anxiety-like behaviors in female adolescent prairie voles

Repeated measures ANOVA was used to determine differences between the dark and light side of a light/dark box, analyzing duration of time spent, distance traveled, and vertical activity in each side, as well as the number of entries into the dark compartment. In female voles, duration of time spent on either side of the light/dark box revealed significant within-subjects differences, *F*(1,33) = 7.174, *p* = .011, η_p_^2^ = .179, indicating CON voles spent more time in the light than the dark while ELSD voles spent more time in the dark than the light (Figure 1A). There were no differences in between-subjects measures, *F*(1,33) = 0.027, *p* = .870 (Figure 1A). The total distance traveled was reduced in female ELSD voles compared to CON, with significant differences found in analyses for both within-subjects, *F*(1,33) = 8.800, *p* = .006, η_p_^2^ = .211, and between-subjects, *F*(1,33) = 8.457, *p* = .006, η_p_^2^ = .204 (Figure 1B). The number of dark of entries female voles performed did not differ based on condition, *t*(33) = 1.644, *p* = .110, (Figure 1C) suggesting both ELSD and CON voles moved between the light and the dark equally. Finally, vertical activity (*i.e.* rearing), was found to be significantly reduced based on within-subjects, *F*(1,33) = 8.384, *p* = .007, η_p_^2^ = .203, and between-subjects, *F*(1,33) = 14.726, *p* = .001, η_p_^2^ = .309, analyses (Figure 1D). Post hoc analyses on duration, distance traveled, and vertical activity were conducted to determine between-subjects effects. Duration in the light was shown to be significantly reduced in female ELSD voles compared to CON voles, *t*(33) = 2.678, *p* = .011, while duration in the dark was significantly increased, *t*(33) = −2.678, *p* = .011 in ELSD female voles. Distance traveled in the light was reduced in female ELSD voles, *t*(33) = 3.853, *p* = .001, but distance traveled in the dark did not differ, *t*(33) = 1.393, *p* = .173 (Figure 1B), indicating female ELSD voles moved less than controls in the light. Female ELSD voles performed less vertical activity in both the light, *t*(33) = 4.426, *p* < .0001, and the dark, *t*(33) = 2.341, *p* = .025 (Figure 1D).

Male prairie voles did not experience the same changes to duration, distance traveled, entries, or vertical activity. Duration of time spent was not impacted by ELSD, with no differences found based on analyses of within-subjects, *F*(1,38) = 0.185, *p* = .670, or between-subjects, *F*(1,38) < 0.001, *p* > .999 (Figure 2A). Analyses of distance traveled indicated a trend, with near-significant reductions of within-subjects measures, *F*(1,38) = 3.582, *p* = .066, η_p_^2^ = .086, but no differences from between-subjects measures, *F*(1,38) = 0.807, *p* = .375 (Figure 2B). ELSD did not impact entries to the dark side of the apparatus, no differences were found based on within-subjects, *F*(1,38) = 0.393, *p* = .534, or between-subjects, *F*(1,38) = 2.100, *p* = .155, analyses (Figure 2C). Similarly, vertical activity was no different in ELSD voles based on analyses of within-subjects, *F*(1,38) = 0.520, *p* .475, and between-subjects, *F*(1,38) = 0.696, *p* = .409, measures (Figure 2D).

**Figure 2.**
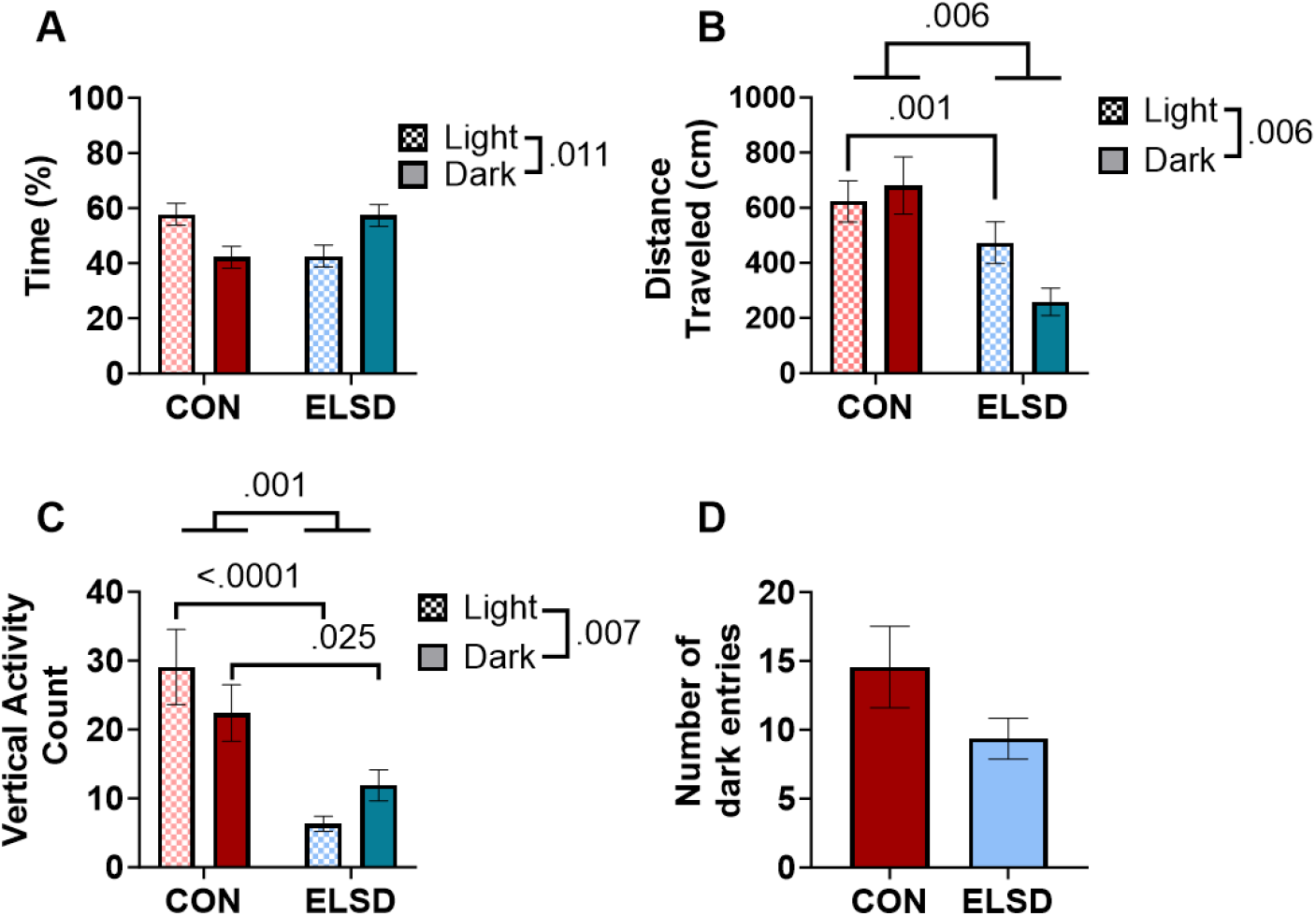
Anxiety-like behavior in female prairie voles. Anxiety-like behavior was quantified with a 10 minute test in the light/dark box. Female ELSD prairie voles showed increased anxiety-like behavior as evidenced by reduced time (**A**), less distance traveled (**B**), and less rearing activity (**C**) in the light compared to Controls. The number of dark entries (**D**) was not significantly different in ELSD females compared to Controls.

**Figure 3.**
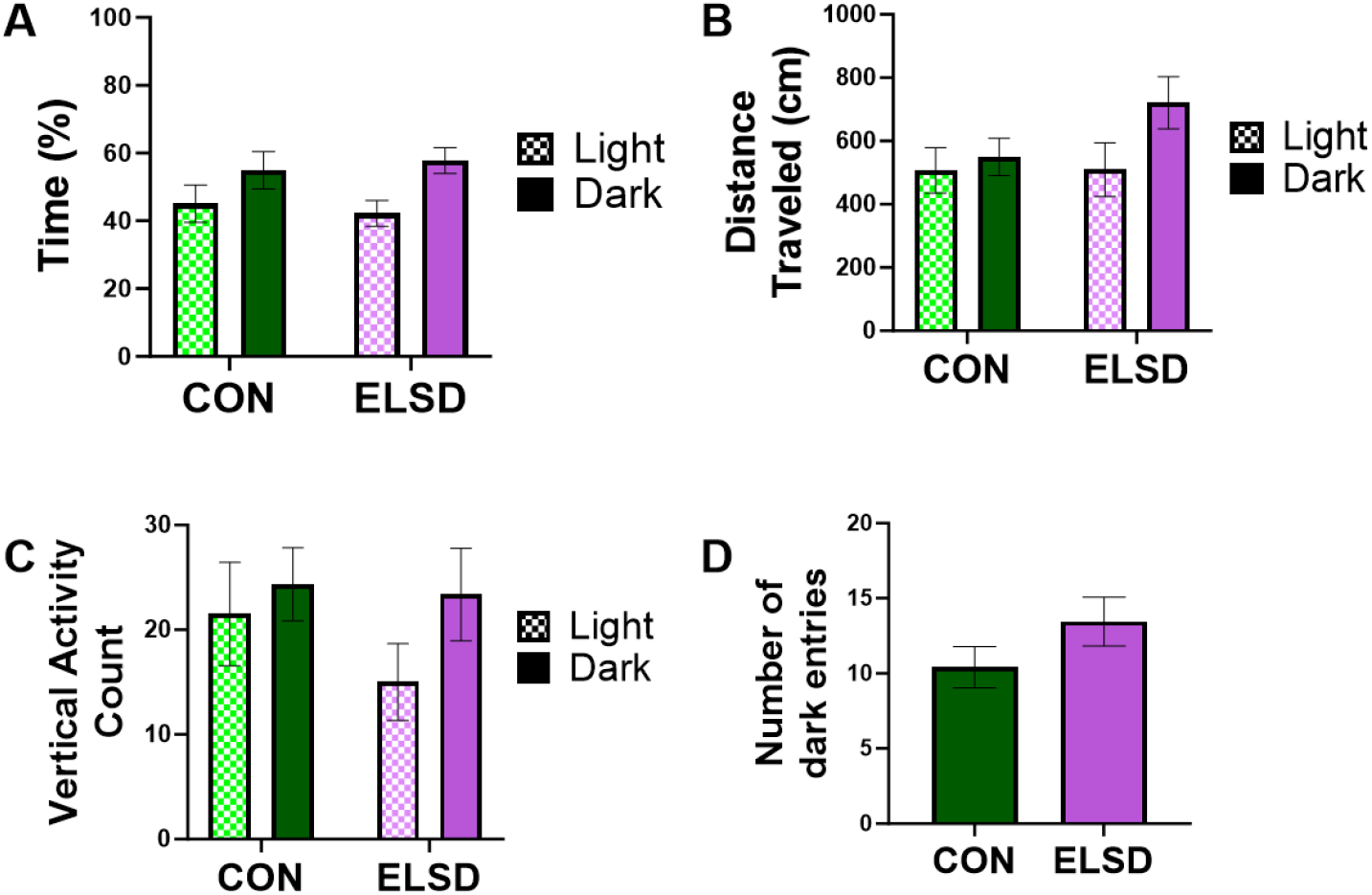
Anxiety-Like behavior in male prairie voles. Male prairie voles did not differ on time (**A**), distance traveled (**B**), rearing activity (**C**), or dark entries (**D**) based on ELSD or CON condition.

**Figure 3.**
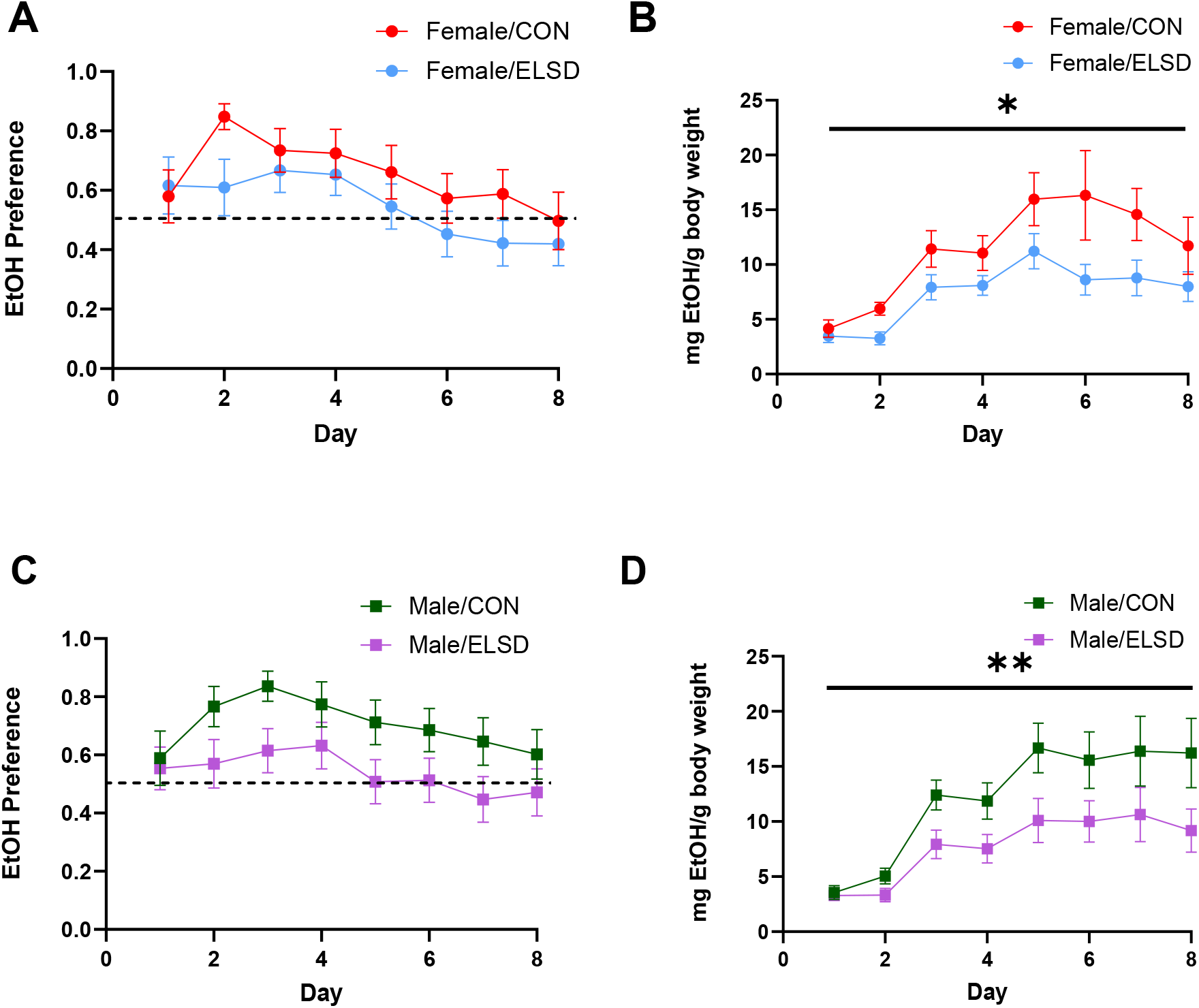
Ethanol preference (left) and consumption (right). Female prairie vole ethanol preference (**A**) was similar between groups, but ethanol consumption was reduced in female ELSD voles compared to controls (**B**). ELSD did not significantly affect male ethanol preference (**C**) or consumption (**D**). Dotted line indicates equal ethanol to water preference, * indicates *p* = .033 and ** indicates *p* = .044.

### Early-life sleep disruption results in reduced ethanol consumption in both males and females

To examine ethanol preference a repeated measures ANOVA was used. Within-subjects analyses showed no differences based on condition between males, *F*(7,252) = 0.632, *p* = .729, and females, *F*(7,210) = 0.891, *p* = .514. Between-subjects analyses revealed a near-significant trend in males, *F*(1,36) = 3.677, *p* = .063, η_p_^2^ = .093, but not in females, *F*(1,30) = 0.498, *p* = .486.

To examine total ethanol consumed, total weight of ethanol consumed (mg) per bodyweight (g) was calculated to normalize ethanol consumption across animals. Within-subjects results were non-significant for both males, *F*(7,245) = 1.577, *p* = .143, and females, *F*(7,203) = 1.101, *p* = .364. Between-subjects analyses revealed a significant effect of ELSD on both males, *F*(1,36) = 4.347, *p* = .044, η_p_^2^ = .108, and females, *F*(1,29) = 5.013, *p* = .033, η_p_^2^ = .147, indicating ELSD results in lower consumption of ethanol compared to CON in both sexes. Together these results reveal no impact of ELSD on ethanol preference, regardless of sex, but ELSD does attenuate total ethanol consumed in both males and females.

### Males that experienced early-life sleep disruption performed less affiliative behavior with their cagemate

Upon the completion of two bottle choice test, a subset of prairie voles were tested for social behavior in a clean home cage with their same sex sibling cage-mate. Affiliative behavior duration was operationally defined as the time paired prairie voles spent huddled and allogrooming. Autogrooming was quantified to determine if any differences in affiliative behavior was related to changes in self-care. A between-subjects ANOVA on affiliative behavior and condition revealed no impact of ELSD on female affiliative behavior, *F*(1,14) = 2.188, *p* = .161 (Figure 4A). Autogrooming duration was no different between ELSD and CON female voles, *F*(1,14) = 0.643, *p* = .436 (Figure 4B). Male ELSD affiliative behavior was significantly reduced in duration compared to CON, *F*(1,21) = 4.743, *p* = .041, η_p_^2^ = .184 (Figure 4C). No differences were found in males regardless of ELSD or CON conditions, *F*(1,21) = 0.130, *p* = .722 (Figure 4D).

**Figure 4.**
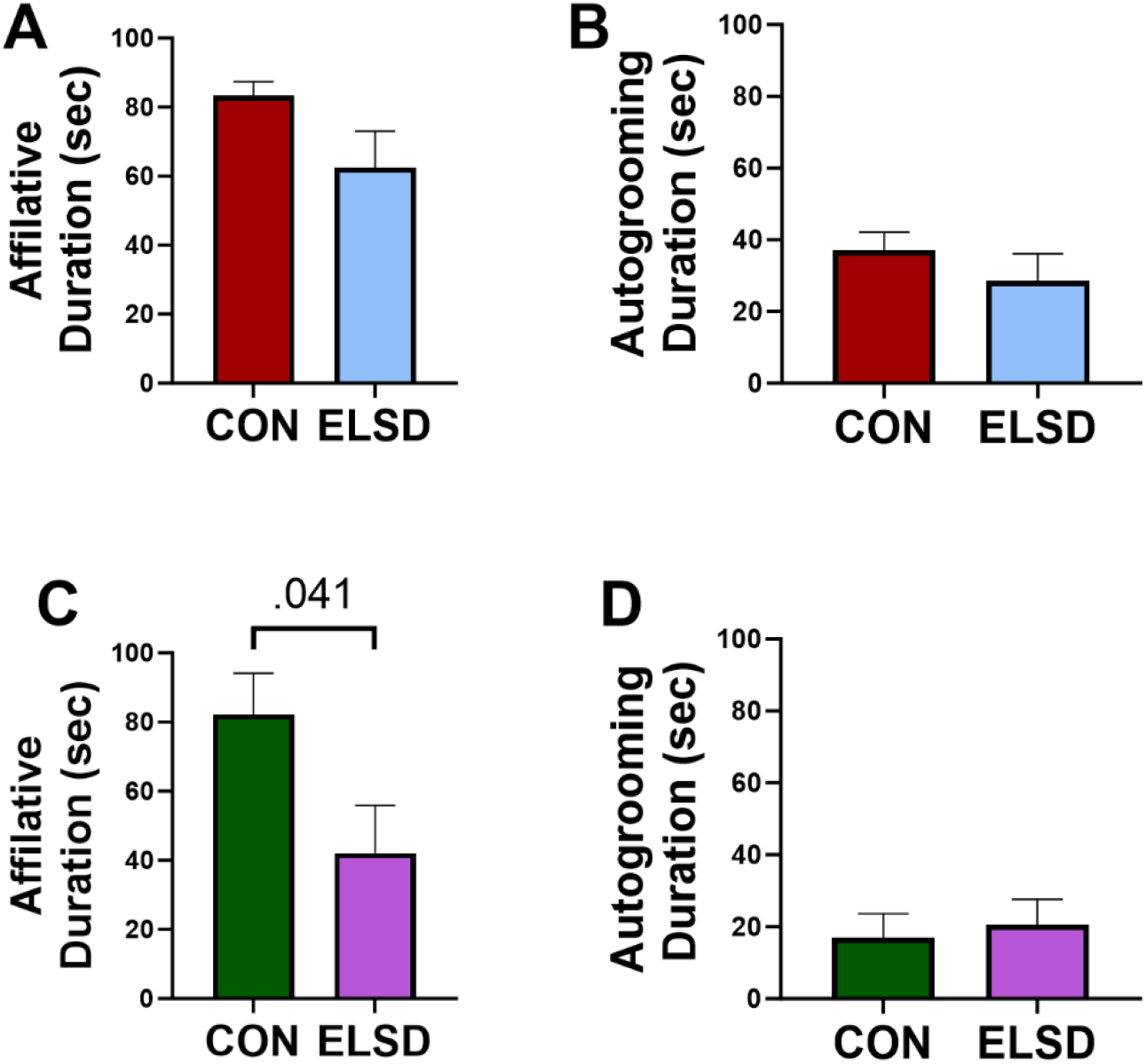
Affiliative behavior duration was defined as the time prairie voles spent huddling and/or grooming their same sex sibling pair. Female affiliative behavior (**A**) and autogrooming (**B**) was similar between CON and ELSD voles. Male ELSD voles had reduced affiliative behavior (**C**) while autogrooming (**D**) remained unchanged.

**Figure 5.**
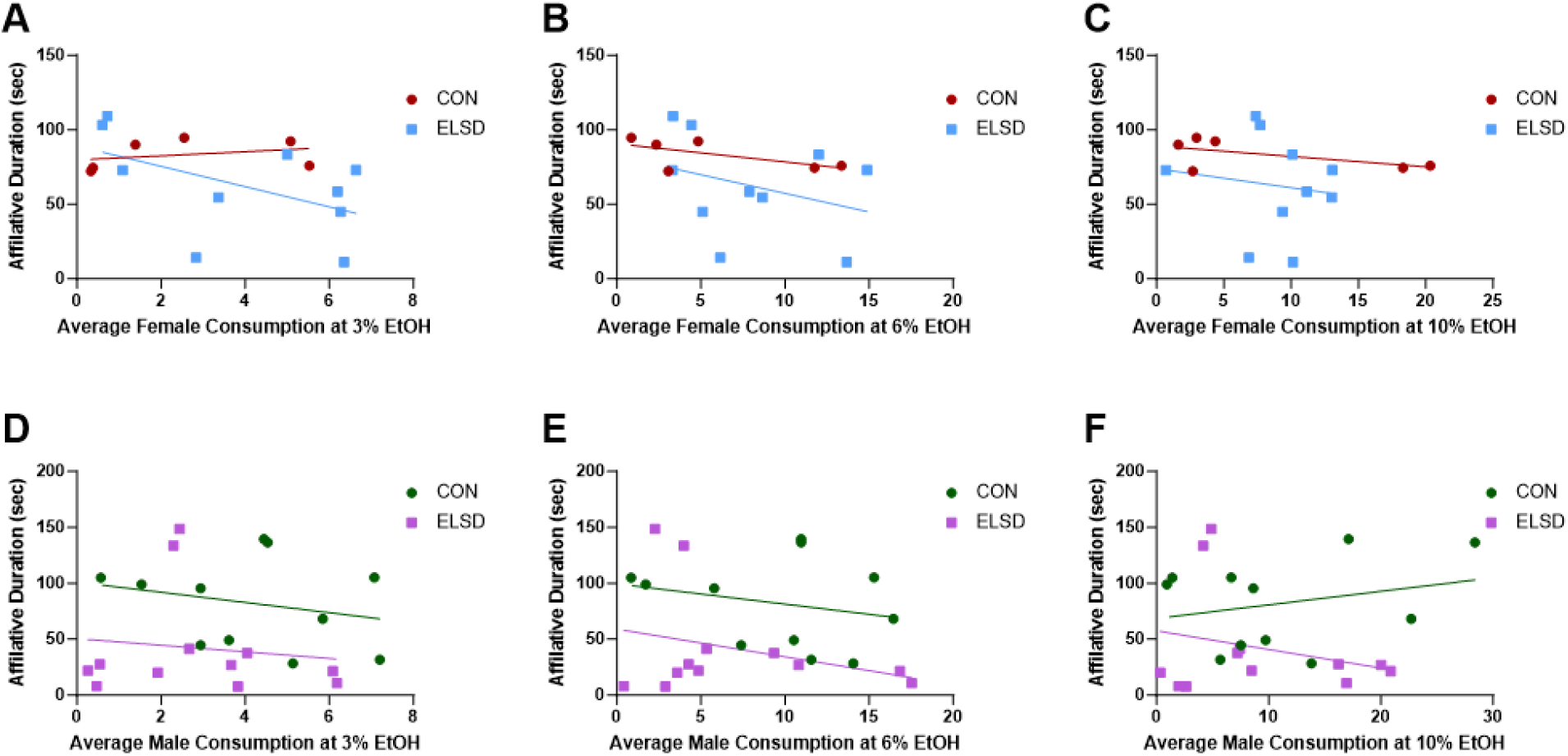
Correlational analyses between affiliative behavior and ethanol consumption yielded no significant relationships between females at 3% (**A**), 6% (**B**), and 10% (**C**) ethanol concentrations. Males also showed no differences at 3% (**D**), 6% (**E**), and 10% (**F**) ethanol concentrations. These data indicate no effect of ethanol consumption on affiliative behavior.

### Ethanol consumption did not impact affiliative behavior

As two-bottle choice tests were performed before affiliative behavior tests, examining the impact of ethanol consumption on affiliative behavior was necessary. Pearson analyses were performed on average ethanol consumed per concentration (*i.e.* 3%, 6%, and 10%) and affiliative behavior. Female CON voles showed no significant correlations between affiliative behavior and 3% ethanol, *r*(6) = .314 *p* = .544, 6% ethanol, *r*(6) = −.628, *p* = .182, or 10% ethanol consumption, *r*(6) = −.594. *p* = .214 (Table 1.1). Male CON voles showed similar results, with no significant relationship between affiliative behavior and 3% ethanol, *r*(11) = -.242, *p* = .473, 6% ethanol, *r*(11) = -.235, *p* = .487, and 10% ethanol, *r*(11) = .262, *p* = .436 (Table 1.1). Female ELSD voles also showed a non-significant relationship between affiliative behavior and 3% ethanol, *r*(10) = -.513, *p* = .129, 6%, *r*(10) = -.323, *p* = .362, and 10% ethanol, *r*(10) = -.140, *p* = .700 (Table 1.2). The relationship male ELSD voles had between affiliative behavior and 3% ethanol, *r*(12) = -.124, *p* = .701, 6% ethanol, *r*(12) = -.292, *p* = .357, and 10% ethanol, *r*(12) = -.256, *p* = .421, also had no significant relationship (Table 1.2). The lack of significance suggests that ethanol consumption did not impact affiliative behavior in both sexes, regardless of condition.

**Table 1.1.** Correlational analyses reveal no significant relationship between affiliative behavior and ethanol consumption in male and female prairie voles.

| CON Females | *M* | *SD* | df | *r* | *p* |
| --- | --- | --- | --- | --- | --- |
| 3% mg/g | 4.706 | 2.773 | 4 | .314 | .544 |
| 6% mg/g | 11.235 | 6.325 | 4 | -.628 | .182 |
| 10% mg/g | 15.010 | 9.38 | 4 | -.594 | .214 |
| CON Males | *M* | *SD* | df | *r* | *p* |
| 3% mg/g | 4.304 | 2.372 | 9 | -.242 | .473 |
| 6% mg/g | 12.137 | 6.252 | 9 | -.235 | .487 |
| 10% mg/g | 16.217 | 11.47 | 9 | .262 | .436 |

**Table 1.2.** Correlational analyses reveal no significant relationship between affiliative behavior and ethanol consumption in male and female prairie voles.

| ELSD Females | *M* | *SD* | df | *r* | *p* |
| --- | --- | --- | --- | --- | --- |
| 3% mg/g | 3.367 | 2.374 | 8 | -.513 | .129 |
| 6% mg/g | 8.007 | 3.782 | 8 | -.323 | .362 |
| 10% mg/g | 9.154 | 5.74 | 8 | -.140 | .700 |
| ELSD Males | *M* | *SD* | df | *r* | *p* |
| 3% mg/g | 3.296 | 2.434 | 10 | -.124 | .701 |
| 6% mg/g | 7.725 | 5.452 | 10 | -.292 | .357 |
| 10% mg/g | 9.979 | 8.37 | 10 | -.256 | .421 |

## DISCUSSION

Early-life sleep can be considered critically important as we have previously shown that adult prairie voles that experienced ELSD display social impairments (Jones et al., 2019), increased ethanol consumption following a stressor (Jones et al., 2020), and impaired cognitive flexibility (Jones et al., 2021). The present study further exemplifies the importance of early-life sleep for adolescent prairie voles as well as highlights sex specific impacts of ELSD. The above data suggests that adolescent female ELSD voles experienced heightened anxiety and both males and females find ethanol to less rewarding.

Furthermore, only males exhibited reduced affiliative behavior, suggesting adolescent male ELSD voles find allogrooming and huddling to be less rewarding compared to controls. When paired with the previous findings, ELSD can be considered to modify behavior across the lifespan, with behaviors in both adolescence and adulthood altered.

The data from the light-dark box highlights sex differences following ELSD in anxiety-like behavior. The current literature shows that there is no difference between unmanipulated male and female mice (Henarejos et al., 2020) or rats (Domonkos et al., 2017) when examining light-dark box behavior, suggesting anxiety-like behavior may be comparable between males and females. Assuming this phenomenon holds true for prairie voles, the differences seen in the present study are not solely related to sex, but an impairment mediated by ELSD. Differences in the excitatory: inhibitory (E:I) ratio in the medial prefrontal cortex (mPFC) may explain these sex specific impacts. We have previously shown that ELSD results in reductions to vesicular glutamatergic transporter 1 expression in the prelimbic region (Jones et al., 2021), suggesting reduced excitation. With this, inhibitory parvalbumin (PV) interneurons have been reported to control anxiety-like behavior in female mice, with increased activity of PV responsible for increased measures of anxiety-like behavior in an open field (Page et al., 2019). The reported change to the E:I balance in the prelimbic region (Jones et al., 2021) allows for increased inhibition, which is likely related to PV activity as they are the population of interneurons with the highest density in the mPFC (Kepecs & Fishell, 2014; Kupferschmidt et al., 2022). Thus, ESLD likely results in increased in PV activity in the mPFC, facilitating the sex specific increased anxiety-like behavior.

The reduced social interaction in male ELSD prairie voles may potentially be related to reductions in reward salience of social interactions. Social interactions have been shown to be highly salient in rats, with rats preferentially lever pressing for interactions with a same sex cage mate over drugs of abuse, such as heroin (Venniro et al., 2019), cocaine (Venniro et al., 2021), and methamphetamine (Venniro et al., 2018). Social interaction in prairie voles has been shown to not only involve dopamine neurotransmission, classically recognized as the reward molecule (Wise, 2004), but prairie voles will also lever press for social interactions. Pharmacological manipulation of dopamine via apomorphine has been shown to facilitate peer partnerships in female prairie voles (Lee & Beery, 2021), highlighting the role of reward circuitry in social behavior. Furthermore, both male and female prairie voles will lever press for access to both an opposite sex and same sex partner (Beery et al., 2021). Together, these studies provide strong evidence for the salience of social interaction. Here, the reduced affiliative behavior in males can be interpreted as a measure of reduced reward seeking behavior in adolescence. Therefore, reduced social interactions can be understood as another measure of reduced reward salience, especially in the highly social prairie vole (Beery et al., 2018).

The sex specific reductions in ethanol consumption may be unique to prairie voles and further highlight the appropriateness of ELSD in prairie voles as a model of human ASD. Similar to humans, prairie voles find ethanol to be highly rewarding (Lewis & June, 1990; Potretzke & Ryabinin, 2019). Here, we report that ELSD results in female prairie voles reducing their ethanol consumption while ethanol preference remains intact. This did not occur in males. Human ASD research has shown that men diagnosed with ASD were more likely to drink alcohol more compared to women (Bowri et al., 2021). This sex specific phenomenon in humans was mimicked here. The mechanism behind this difference in ethanol consumption may be related to changes in either dopamine or oxytocin neurotransmission. Studies have shown that individuals with ASD have shown reductions in dopamine receptor D2 mRNA (Brandenburg et al., 2020) and single nucleotide polymorphisms in the *oxytocin receptor* gene (Wermter et al., 2010). Changes to these systems likely alters both social behavior and reward seeking behavior, leading to the reductions in seen in the human population regarding socialization and reward seeking (Baumester et al., 2023). ELSD may result in similar impacts on these receptors, resulting in the behavioral changes seen here.

## CONCLUSIONS

Here we report that ELSD results in sex specific changes to behavior. Female prairie voles experience heightened anxiety and reduced ethanol consumption while reduced social interactions with siblings is the only impacted behavior in male prairie voles. As such, paired with our previous works (see Jones et al., 2019; 2020; 2021), we present further evidence to support both face and construct validity of using ELSD and prairie voles to model human ASD. More research is required to further understand the behavioral findings here. Close examination of receptors implicated in ASD (*i.e.* dopamine D2 and oxytocin) using our ELSD model would provide a mechanism to explain the observed behavior at this age. Additionally, examination of brain tissue either during or immediately following ELSD may provide some insight to differences in signaling molecules that are involved with neuronal development and maturation during pre-weaning ages (*i.e.* P14-P21).

## Acknowledgements

This work was supported by VA Biomedical Laboratory Research & Development (BLR&D) Career Development Award (CDA) # IK2 BX002712, Portland VA Research Foundation, Brain & Behavior Foundation NARSAD Award, Collins Medical Trust, and NIH EXITO Institutional Core, #UL1GM118964 to MML; NIH T32 5T32AA7468-29 and NIH T32 5T32HL083808-10 to CET and NIH T32 AA007468-36 to DEG. These contents do not represent the views of the U.S. Department of Veterans Affairs or the United States Government.

